# Distributed Bayesian Networks Reconstruction on the Whole Genome Scale

**DOI:** 10.1101/016683

**Authors:** Alina Frolova, Bartek Wilczynski

## Abstract

**Background:** Bayesian networks are directed acyclic graphical models widely used to represent the probabilistic relationships between random variables. They have been applied in various biological contexts, including gene regulatory networks and protein-protein interactions inference. Generally, learning Bayesian networks from experimental data is NP-hard, leading to widespread use of heuristic search methods giving suboptimal results. However, in cases when the acyclicity of the graph can be externally ensured, it is possible to find the optimal network in polynomial time. While our previously developed tool BNFinder implements polynomial time algorithm, reconstructing networks with the large amount of experimental data still leads to computations on single CPU growing exceedingly.

**Results:** In the present paper we propose parallelized algorithm designed for multi-core and distributed systems and its implementation in the improved version of BNFinder - tool for learning optimal Bayesian networks. The new algorithm has been tested on different simulated and experimental datasets showing that it has much better efficiency of parallelization than the previous version. BNFinder gives comparable results in terms of accuracy with respect to current state-of-the-art inference methods, giving significant advantage in cases when external information such as regulators list or prior edge probability can be introduced.

**Conclusions:** We show that the new method can be used to reconstruct networks in the size range of thousands of genes making it practically applicable to whole genome datasets of prokaryotic systems and large components of eukaryotic genomes. Our benchmarking results on realistic datasets indicate that the tool should be useful to wide audience of researchers interested in discovering dependencies in their large-scale transcriptomic datasets.

## Background

Bayesian networks (BNs) are graphical representations of multivariate joint probability distributions factorized consistently with the dependency structure among variables. In practice, this often gives concise structures that are easy to interpret even for non-specialists. A BN is a directed acyclic graph with nodes representing random variables and edges representing conditional dependencies between the nodes. Nodes that are not connected represent variables that are independent conditionally on their parent variables [1]. In general, inferring BN structure is NP-hard [2], however it was shown by Dojer [3] that it is possible to find the optimal network structure in polynomial time when datasets are fixed in size and the acyclicity of the graph is pre-determined by external constraints. The latter is true when dealing with dynamic BNs or when user defines the regulation hierarchy restricting the set of possible edges in case of static BNs. This algorithm was implemented in BNFinder - a tool for BNs reconstruction from experimental data [4].

One of the common use of BNs in bioinformatics is inference of interactions between genes [5] and proteins [6]. Even though it was originally developed for this purpose, BNFinder is a generic tool for reconstructing regulatory interactions. Since its original publication, it was successfully applied to linking expression data with sequence motif information [7], identifying histone modifications connected to enhancer activity [8] and to predicting gene expression profiles of tissue-specific genes [9]. Even though it can be applied to many different datasets, the practical usage of the algorithm is limited by its running times that can be relatively long. Since the algorithm published by Dojer [3] has a capacity to be parallelized by design and the current version of BNFinder [10] has only a simple parallelization implemented, we have developed a new version that takes advantage of multiple cores via the python multiprocessing module and gives better performance.

## Implementation

The general scheme of the learning algorithm is the following: for each of the random variables find the best possible set of parent variables by considering them in a carefully chosen order of increasing cost function. Current parallelization in BNFinder version 2 [10] can be considered **variable-wise** as it distributes the work done on each variable between the different threads. However, such approach has natural limitations. Firstly, the number of parallelized tasks cannot exceed the number of random variables in the problem, meaning that in the cases where only a few variables are considered (e.g. in classification by BNs) we get a very limited performance boost. Secondly, **variable-wise** parallelization might be vulnerable (in terms of performance) to the datasets with highly heterogeneous variables, i.e. variables whose true dependency graph has a wide range of connections. As the time spent on computing parent sets for different variables varies - it leads to uneven load of threads. In biology we usually observe networks with scale-free topology consisting of a few hub nodes with many parents and a large number of nodes that have one or small number of connections [11]. If one applies **variable-wise** algorithm to such networks the potential gain in the algorithm performance is not greater than in the case where all the nodes have as many parents as the hub node with the largest parent set.

While **variable-wise** algorithm is the most straightforward one, it is also possible to consider different possible parents sets in parallel denoting **set-wise** algorithm. It means that in the first step we compute singleton parents sets using all available threads, in the second step we compute two-element parents sets in parallel and so on, until we reach parents sets size limit or score function limit. However, **set-wise** algorithm requires more synchronizations between the threads [12] in comparison with **variable-wise**. On top of that allocating large number of cores to the variable whose parents set is very quick to compute may result in lower performance due to context switching. As it is difficult to tell, which problem might be more important in practice, we have implemented and tested two approaches: **set-wise** only and **hybrid** one - a combination of **variable-wise** and **set-wise**.

Figures 1,2 and 3 show python pseudocode for **variable-wise**, **set-wise** and **hybrid** algorithms accordingly, which was simplified in comparison to the original implementation for better illustration. As was stated above **set-wise** algorithm uses each given core to compute parents sets for one gene and after finding parents it proceeds with the next gene. On the contrary **hybrid** algorithm uniformly distributes cores between genes, for example, if user has 3 genes in the network and 6 cores available, each gene will have 2 cores for computing its parents set. If there are 7 cores available, one gene will have 3 cores, while other two genes - 2 cores. Thus, once the gene is processed the freed cores cannot be allocated to other genes, which may be a potential disadvantage.

**Figure 1.**
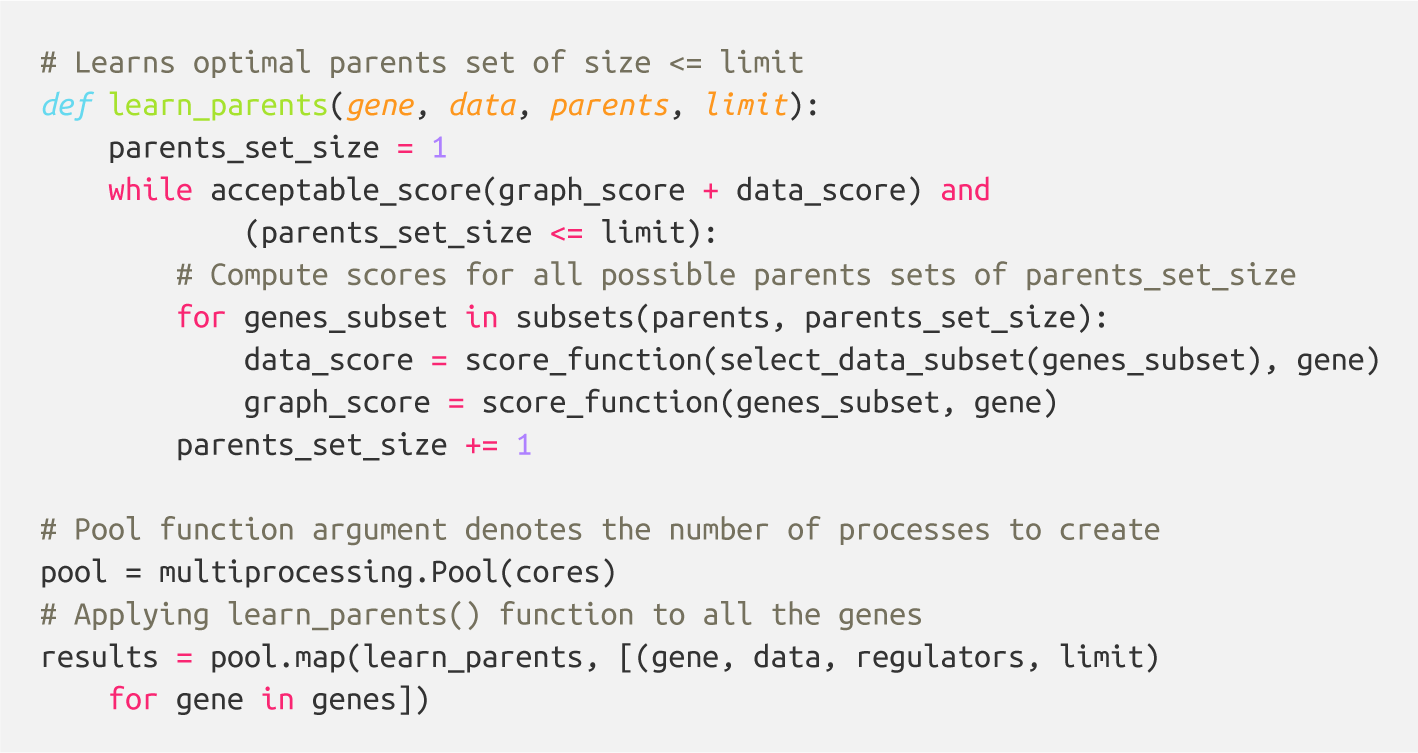
Variable-wise algorithm python pseudocode.

**Figure 2.**
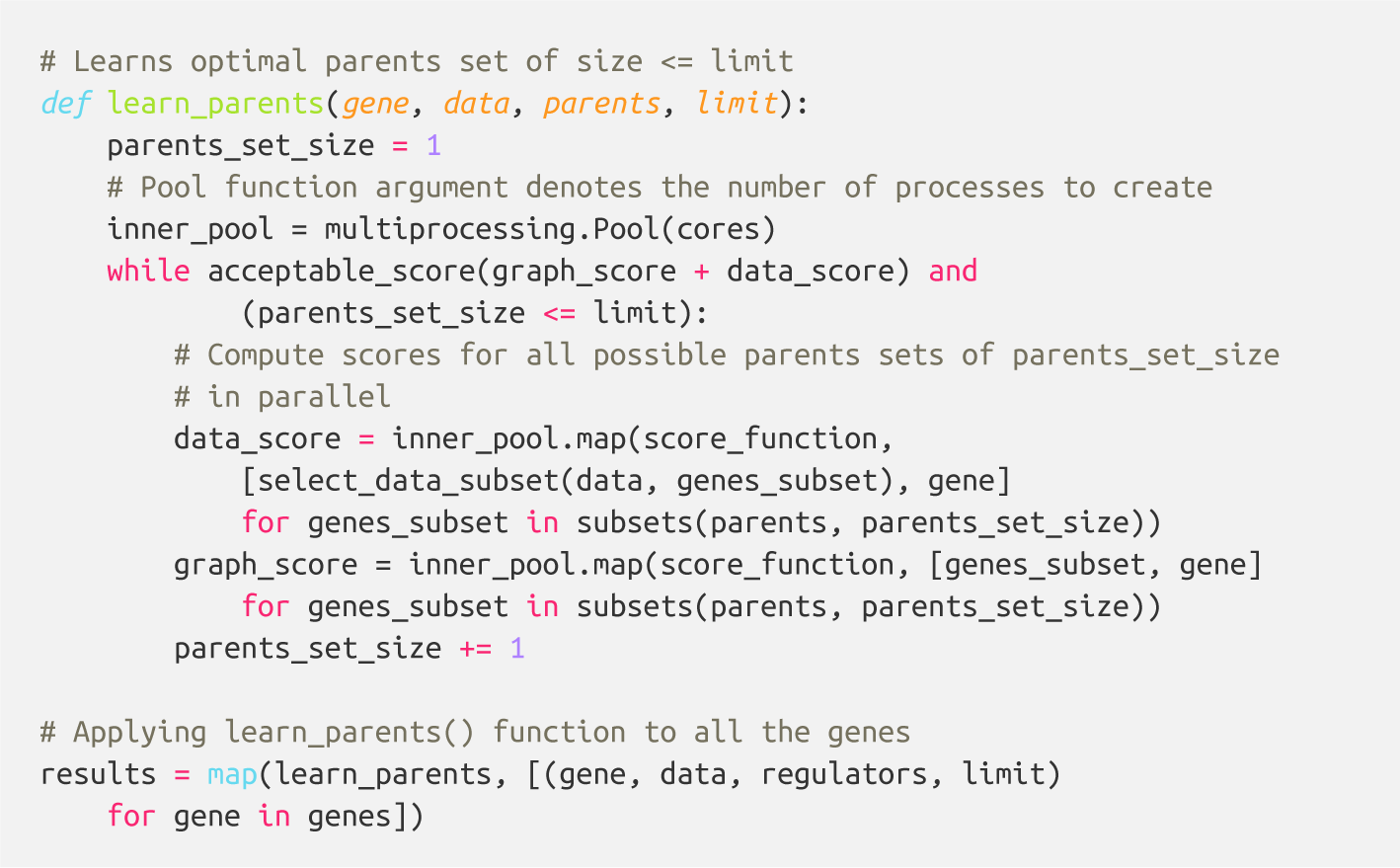
Set-wise algorithm python pseudocode.

**Figure 3.**
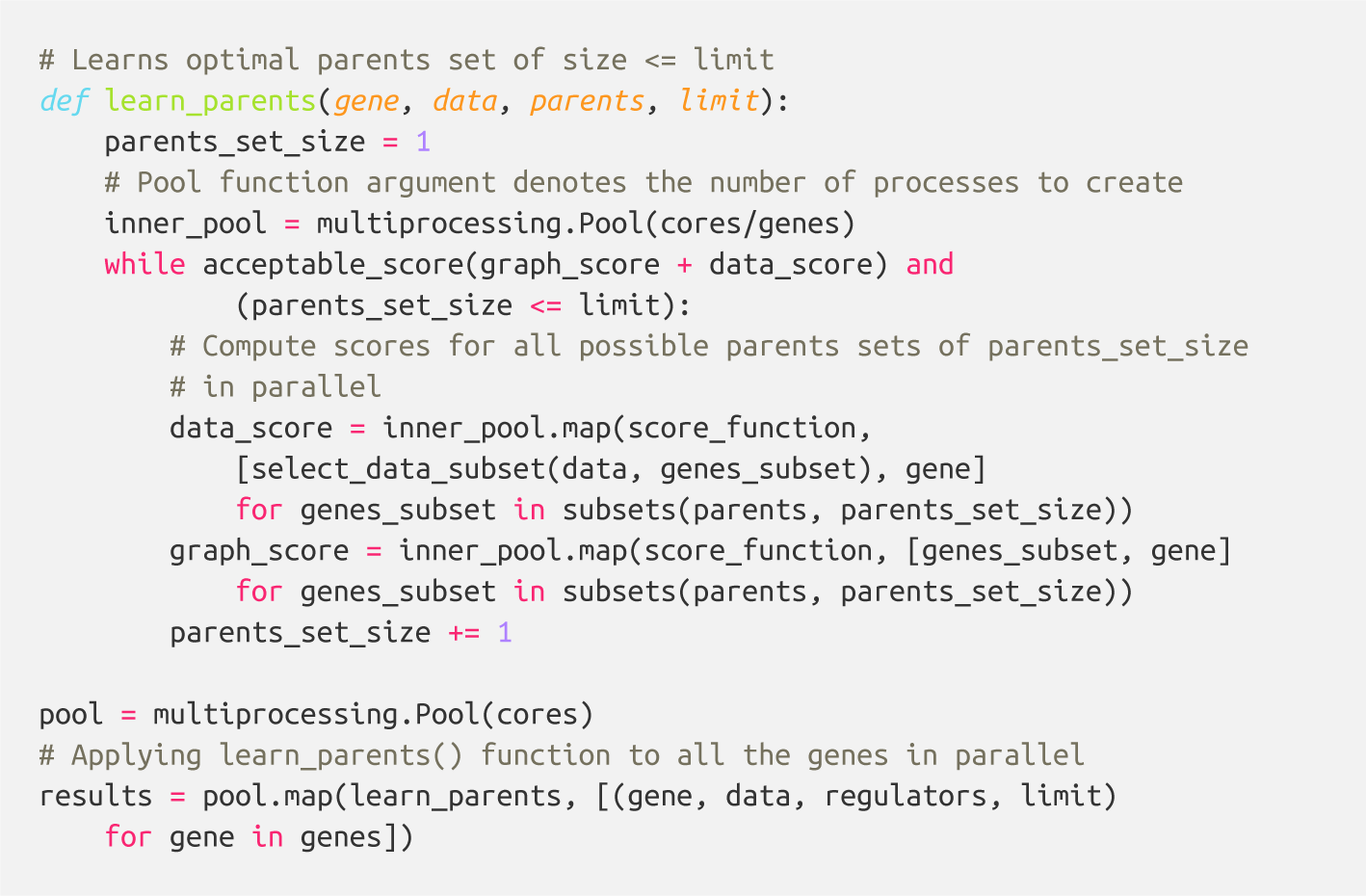
Hybrid algorithm python pseudocode.

So, the pure theoretical complexity of **set-wise** (left side of inequality) and **hybrid** (right side of inequality) algorithms can be described in following way:

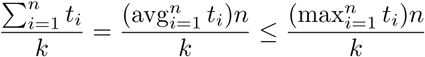

where *k* is the cores number, *n* is the number of random variables, and *t*_*i*_ is the time one needs to compute optimal parents set for the *i*^*th*^ variable using one core. Thus, the time to reconstruct the whole network in case of **set-wise** approach is the sum of time needed for each random variable, which is in fact average time one spends on finding the parents set for one variable, while inferring BN with **hybrid** approach is bounded by the maximum time one spends on one variable.

## Results

### Performance testing

#### Algorithms comparison

We compared implementations of three different algorithms: **variable-wise**, **set-wise** and **hybrid**. The original implementation (**variable-wise**) serves as a baseline for computing the speedup and efficiency of the parallelization. For testing we used synthetic benchmark data as well as real datasets concerning protein phosphorylation network published by Sachs et al. [13]. The efficiency is defined as speedup divided by the number of cores used.

**Set-wise** and **hybrid** algorithms performance on 20 genes synthetic network was very similar, while the speedup and efficiency comparison revealed more differences between the algorithms (See Figure 4). There is no regulators list for this network, making BNFinder to reconstruct Dynamic Bayesian network. **Hybrid** algorithm showed more unstable behavior, performing better when the number of cores correlates with the number of genes. It outperformed **set-wise** when the number of cores exceeded the number of genes two times at least, however it didn’t show any speedup when increasing the number of cores from 42 to 50. The latter is easily explained by the algorithm design, since running time is bound by the most computationally complex variable, using 41-59 cores cannot give performance boost as long as this variable provided with one core only.

**Figure 4.**
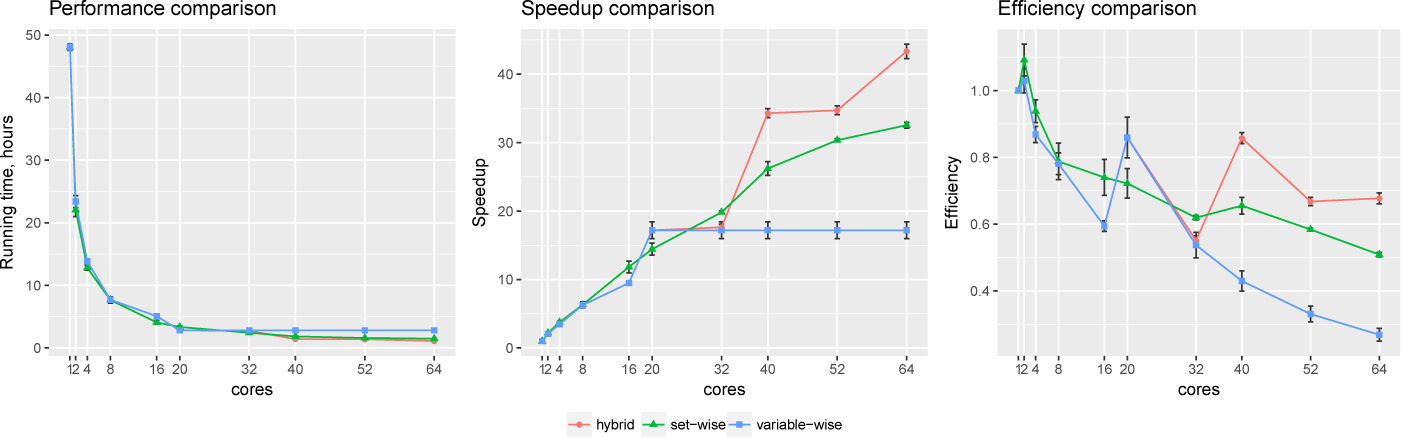
Synthetic data testing. Comparing performance, speedup and efficiency of algorithms on synthetic benchmark data: 20 variables x 2000 observations.

The Sachs et al. network we tried next has 11 proteins, the regulators are selected from those proteins and introduced in the cascade manner, which denotes expected layer structure of the signaling pathway. First layer can regulate all the following, while each next one cannot regulate previous layers. On the first layer only singleton parents set is possible consisting of *plcg* protein, on the second layer we have two regulators, *PIP3* and previously defined *plcg*, making it possible to search for singleton and two-element parents sets for the rest of proteins, and so on.

Sachs et al. data showed significant difference between two algorithms. Clearly, the **set-wise** algorithm outperforms the **hybrid** one: using 11 cores it showed 8x speedup, while the **hybrid** algorithm showed only 1.5x speedup (See Figure 5). **Hybrid** algorithm performance was hindered by highly heterogeneous variables in the input data, because out of the 11 proteins in the network *pakts473* has 6 parents, *p44/42* has 3 parents, while others have 1-2 parents. Importantly, the better performing algorithm is also the one showing more consistent behavior.

**Figure 5.**
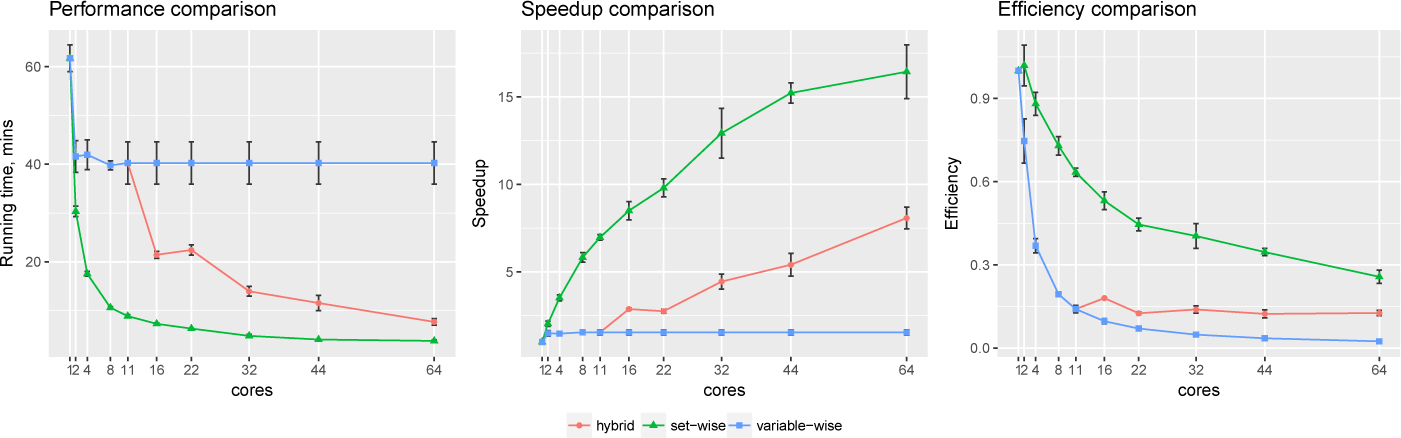
Sachs et al. data testing. Comparing performance, speedup and efficiency of algorithms on protein phosphorylation data: 11 variables x 1023 observations [13].

However, the way how regulators are introduced in the input file produces uneven load by itself, as each next variable has bigger set of potential parents. Therefore, we generated more benchmark data with different number of regulator and target genes, where we could define regulators in one layer manner (one line list) or similar to Sachs data. Moreover, underlying structure of generated networks was designed to be heterogeneous. Namely, it contains genes with gradually increasing number of parents: first gene has zero regulators, second gene - one regulator, third - two regulators and so on. The datasets were generated with *BayesGen.py* script that is included in the supplementary material. It takes the desired connectivity between variables and simulates the observations as emissions from a Bayesian Network with bimodal Gaussian distributions of variables.

The results of multiple tests showed that introducing complex layer structure of regulators always resulted in **hybrid** algorithm poor performance. It either showed much worse results regardless of the number of cores as on Figure 5 or it showed comparable speedup when number of cores was three times bigger than number of genes. In cases when regulators were supplied as one single list both algorithms showed results similar to Figure 4, namely **set-wise** algorithm was better when number of cores was less than number of genes, while **hybrid** one was better in the opposite case (although there was no such dramatical difference as in case of layered regulators structure). The more observations per regulator-target interaction we had, the better BNFinder predicted the network structure. However, as we studied running times per gene we observed that variables with biggest number of parents not always resulted in longest computations. The latter is explained by how the scoring function works, BNFinder stops when the penalty for increasing the set of parents is so big that it cannot improve beyond what it has already found. In general, if the optimal parent set is very good in predicting the child variable value BNFinder will finish searching earlier. It means that the whole family of three-element parents set can have worse score than two-element parents set, but the algorithm will proceed further because it has not reached penalty on increasing the set size yet.

In particular, running times also depend on the number of observation and number of nodes in the networks. For example, Figure 6 shows that increasing the number of observation by 10x leads to 12x longer running time for the network of the same size, while 8 genes network takes 3 times longer to compute in comparison to 7 genes network having the same number of observations.

**Figure 6.**
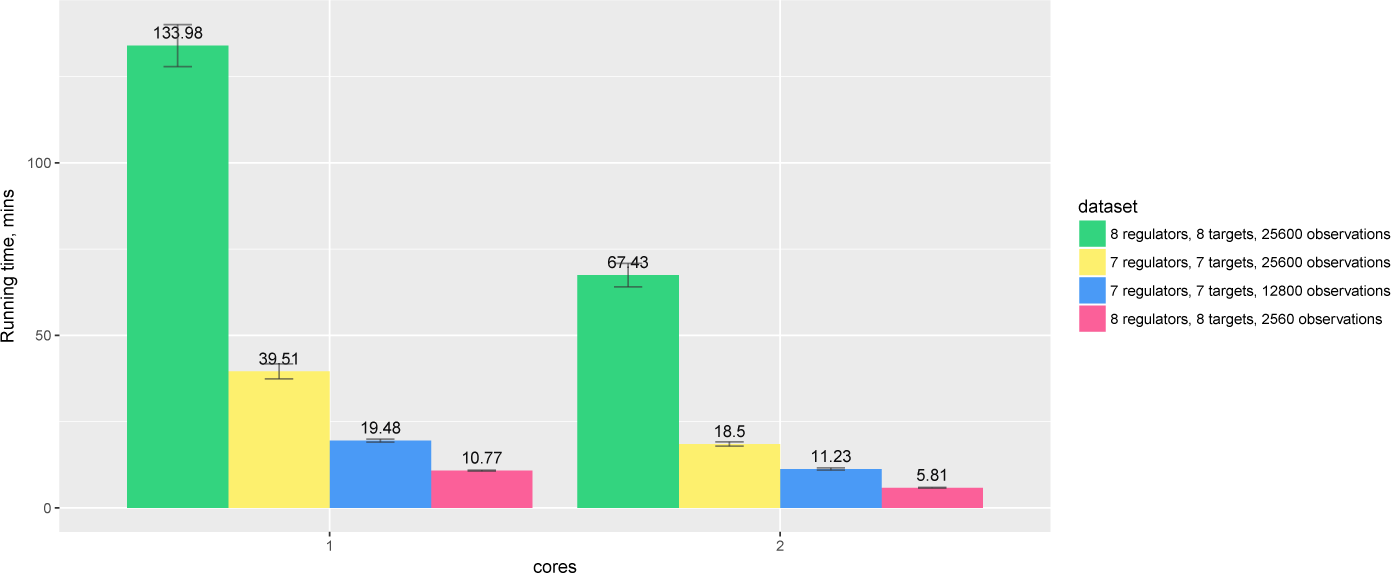
Hybrid algorithm performance on the datasets of difference size. The tendency is preserved for larger number of cores as well.

Since there is no obvious winner between **set-wise** and **hybrid** algorithms, we decided to provide users with both options with **set-wise** being default algorithm. Tests on benchmark and Sachs data were performed on the same server with AMD Opteron (TM) Processor 6272 (4 CPUs with total of 64 cores) and 512GB RAM. During the tests server was loaded only by regular system processes, but to ensure statistical significance we performed each test five times, plotting average values with standard deviations.

#### Distributed computations testing

For the new version of BNFinder we also implemented an option for distributed usage of the tool. The idea is quite simple and did not require specific python libraries or tools. The user has to submit the file with subset of genes as input argument, so BNFinder can calculate partial result. When all the genes are processed user must place all the results into one folder and run BNFinder again, so it will aggregate the results.

For the tests we chose Challenge 5 (Genome-Scale Network Inference) data from DREAM2 competition (Dialogue for Reverse Engineering Assessments and Methods) [14, 15]. Challenge data is a log-normalized compendium of Escherichia coli expression profiles, which was provided by Tim Gardner to DREAM initiative [16]. The participants were not informed about the data origin and were provided only with 3456 genes x 300 experiments dataset and the list of transcription factors.

BNFinder was tested with different parents set limit parameter value (i.e. maximum number of potential parents), which increases the computation time dramatically in non-linear way, especially in case of dataset with many variables. We compared **set-wise** algorithm performance with context likelihood of relatedness (CLR) algorithm - an extension of the relevance networks approach, that utilizes the concept of mutual information [16]. We chose CLR, because it is very fast and easy to use tool, which provides good results. In addition, CLR-based algorithm - synergy augmented CLR (SA-CLR) was best performed algorithm on Challenge 5 [17].

The CLR tests were performed on the GP-DREAM platform, designed for the application and development of network inference and consensus methods [18]. BNFinder tests were done within Ukrainian Grid Infrastructure [19], which was accessed through nordugrid-arc middleware (arc client version 4.1.0 [20]), so the tasks submitting process was automated and unified. The results in Table 1 are not precisely comparable due to differences in used hardware, especially when using such heterogeneous environment as Grid. In addition, we could not obtain stable number of cores over time with the Grid, as the clusters were loaded with other tasks. However, running times can give rough estimate for those who plan to use BNFinder on large datasets.

**Table 1.** DREAM2 Challenge 5 data testing. CLR with cutoff means limiting output results to 100000 genes interactions. *l* stands for BNFinder parents sets limit

|  | CLR | CLR with cutoff | BNF, *l*=1 | BNF, *l*=2 | BNF, *l*=3 |
| --- | --- | --- | --- | --- | --- |
| CPU time, hours | 0.7879 | 0.1999 | 2.0021 | 383.7149 | 109200 |
| Actual time, hours | 0.2626 | 0.0666 | 0.0667 | 12.7904 | 336 |
| CPU number | 3 | 3 | 30 | 30 | ~325 |

Even though computing with parents sets limit 3 takes significant amount of time and resources, it is clear that BNFinder is able to reconstruct genome-scale datasets, significantly broadening its application range after it was adapted to parallel and distributed computing.

### Accuracy testing

Previously we compared accuracy of BNFinder algorithm with Banjo [4] on data provided with the tool and separately on Sachs data [10], which we used in this work to test the performance. Here we performed accuracy testing on 14 different datasets, both synthetic and taken from microarray experiments.

#### DREAM2 Genome Scale Network Inference

3456 genes x 300 experiments dataset, log-normalized compendium of Escherichia coli expression profiles described above [14]. Transcription factors list is provided with the data.

#### DREAM4 In Silico Network Challenge

time course datasets showing how the simulated network responds to a perturbation and how it relaxes upon its removal. There are 5 different datasets with 10 and 100 genes each. For networks of size 10, datasets consist of 5 different time series replicates, while networks of size 100 has 10 time series replicates. Each time series has 21 time points [21].

#### Yeast time series

102 genes x 582 experiments datasets with time series after drug perturbation from the yeastrapamycin experiment described in Yeung et al [22]. There are 582/6 = 97 replicates (the 95 segregants plus two parental strains of the segregants), each with measurements at 6 time points. Prior probabilities of genes regulations are provided.

#### Yeast static

85 genes x 111 experiments subset of the data used for network inference in yeast by Brem et al. [23]. Prior probabilities of genes regulations are provided. TF-gene regulations were extracted from YEASTRACT repository (http://www.yeastract.com, version 2013927).

#### Yeast static synthetic

2000 genes x 2000 experiments dataset generated using GNW simulator [24] by extracting parts of known real network structures capturing several of their important structural properties. To produce gene expression data, the simulator relies on a system of non-linear ordinary differential equations. TF-gene regulations were extracted from YEASTRACT repository (version 2013927). The adjacency matrix of true underlying network structure of this dataset has symmetrical form, therefore it is not possible to evaluate the direction of interaction in this case.

DREAM2 data was downloaded from the challenge website, DREAM4, YeastTS (time series), and Brem data was imported from NetworkBMA [26] R package, while synthetic Yeast static data (GNW2000) was imported from NetBenchmark [25] R package.

Table 2 summarizes gene regulatory network inference methods we used for benchmarking. Specifically, we used FastBMA method implemented in NetworkBMA R package, while the rest of the methods were accessed through NetBenchmark, a bioconductor package for reproducible benchmarks of gene regulatory network inference. The methods were used with default parameters, while for FastBMA, Genie3 and BNFinder prior edge probabilities and TF-gene regulations were supplied where applicable.

**Table 2.** Gene regulatory network inference methods used for benchmarking.

| Inference Approach | Method |
| --- | --- |
| Co-expression algorithms | MutRank [28] |
| Information-theoretic approaches | CLR [16], ARACNE [29], PCIT [30], C3NET [31] |
| Feature selection approaches | MRNET [32], MRNETB [33], Genie3 [34] |
| Bayesian model averaging | FastBMA [35] |

We used two metrics two asses methods performance: Area Under the Precision Recall curve (AUPR) or Area Under Receiver Operating Characteristic curve (AU-ROC), implemented in MINET R package [27]. However, this approach gives an estimation of the global behavior of the method, therefore in NetBenchmark package Bellot at al. evaluated the inferred networks using only the top best 20% of the total number of possible connections [25]. The latter allows to correctly compare methods with sparse and concise outputs. We used both MINET and NetBenchmark evaluation functions in order to assess the impact on methods rankings.

FastBMA, Genie3 and BNFinder methods allow to infer directed interactions, therefore they were additionally evaluated on directed gold networks (where applicable), while for the undirected evaluation gold network adjacency matrices were converted to symmetrical ones (higher edge probabilities are preserved) as well as outputs of directed methods.

In case of static gene expression data FastBMA can infer the regulators of a particular gene by regressing it on the expression levels of the other genes. Therefore we used the method with time series data only.

We mostly used DREAM2 data for running times tests and evaluated accuracy of CLR method only. The results of BNFinder and CLR tests are shown in Table 3.

**Table 3.**
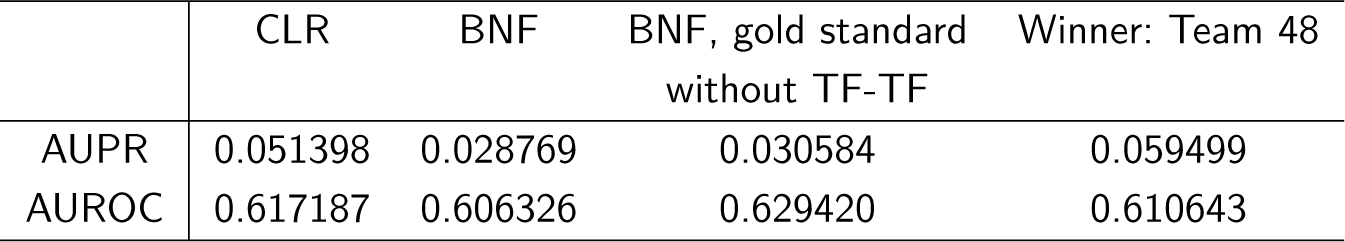
DREAM2 Genome Scale Network Inference. BNFinder and CLR are compared with the best scored method using 100% of output interactions. BNFinder is used with parents sets limit 1 and suboptimal parents sets 100, CLR is used with default parameters. Evaluation of BNF on TF-TF free gold standard is provided as well. All the output interactions are considered for calculating areas under curves.

We ranked AUROC and AUPR values across all the methods for each of five 10 and 100 genes network from DREAM4 challenge. Using different evaluation strategies for 10 genes networks showed quite a different results (Figure 7, 8), while 100 genes network results were more consistent among MINET and NetBechmark (Figure 9, 10). We believe that networks of a small size might not be a good benchmark data as even a slightest change in the obtained scores might disrupt rankings dramatically. This is especially valid for the methods, which output might differ each run due to the nature of underlying algorithms (e.g. regression, greedy hill climbing). There is no single best method for DREAM4 data, while Genie3, MRNET, MRNETB, FastBMA, and BNFinder scored first at least once.

**Figure 7.**
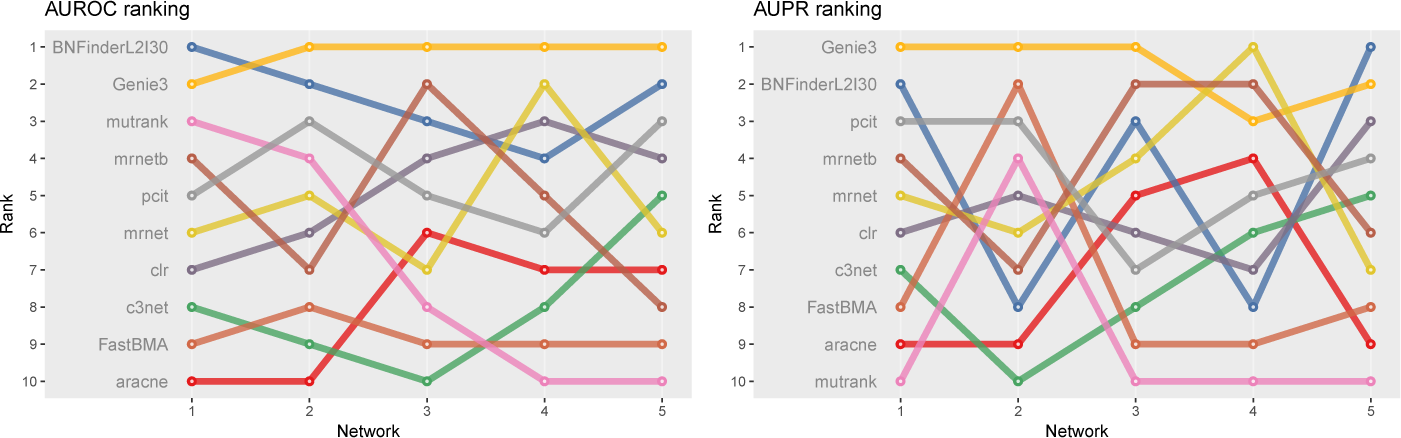
DREAM4 10 genes network, evaluation by MINET package. Area under ROC and PR curves are ranked across different methods. BNFinder is used with parents sets limit 2 and suboptimal parents sets 30.

**Figure 8.**
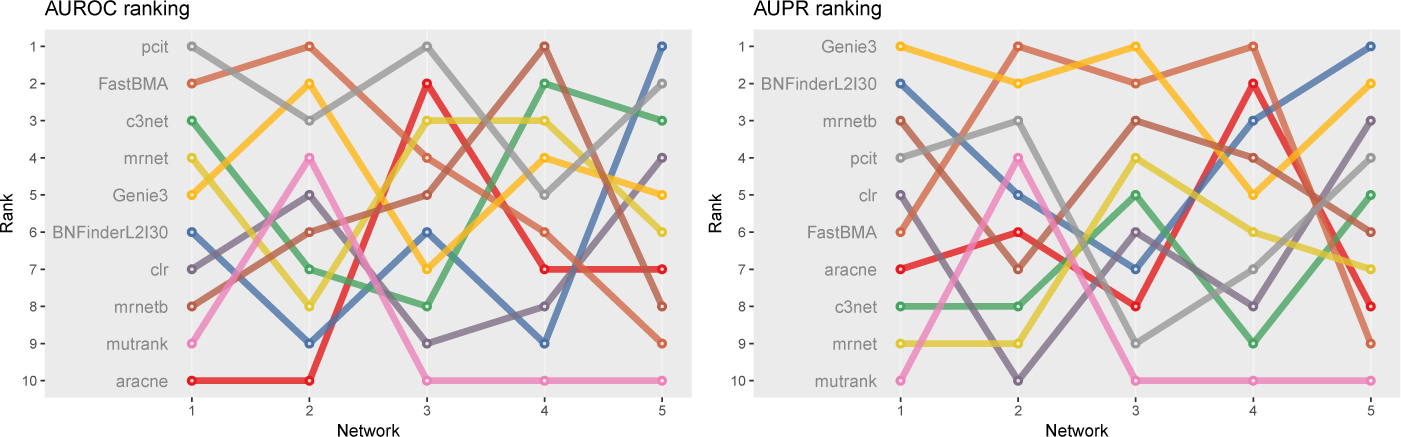
DREAM4 10 genes network, evaluation by NetBenchmark package. Area under ROC and PR curves are ranked across different methods. BNFinder is used with parents sets limit 2 and suboptimal parents sets 30.

**Figure 9.**
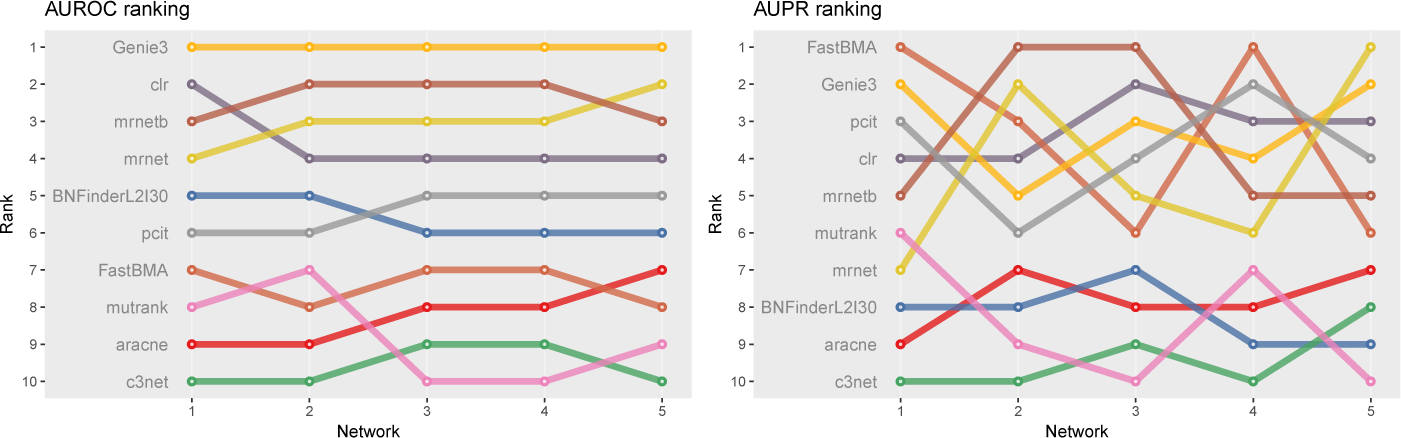
DREAM4 100 genes network, evaluation by MINET package. Area under ROC and PR curves are ranked across different methods. BNFinder is used with parents sets limit 2 and suboptimal parents sets 30.

**Figure 10.**
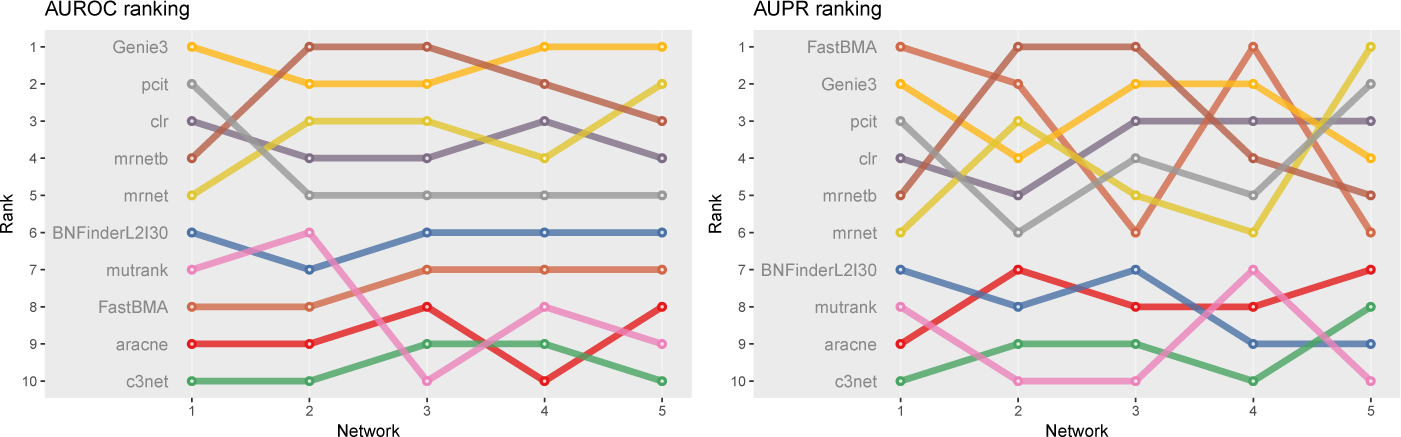
DREAM4 100 genes network, evaluation by NetBenchmark package. Area under ROC and PR curves are ranked across different methods. BNFinder is used with parents sets limit 2 and suboptimal parents sets 30.

Yeast time series network inference showed extremely bad results for all the methods with MutRank having slightly better AUROC=0.58 and AUPR=0.07 values according to MINET package. In contrast to synthetic DREAM4 data which has 21 time points, YeastTS has only 6, which could explain worse results.

Surprisingly, BNFinder significantly outperformed other methods when reconstructing network from Brem at al. static gene expression data (Figure 11). Importantly, Genie3 method was also supplied with the same regulators list as BNFinder, but it has led to worse results contrary to using Genie3 without regulators. The other methods were used without any additional prior information as implemented in NetBenchmark package.

**Figure 11.**
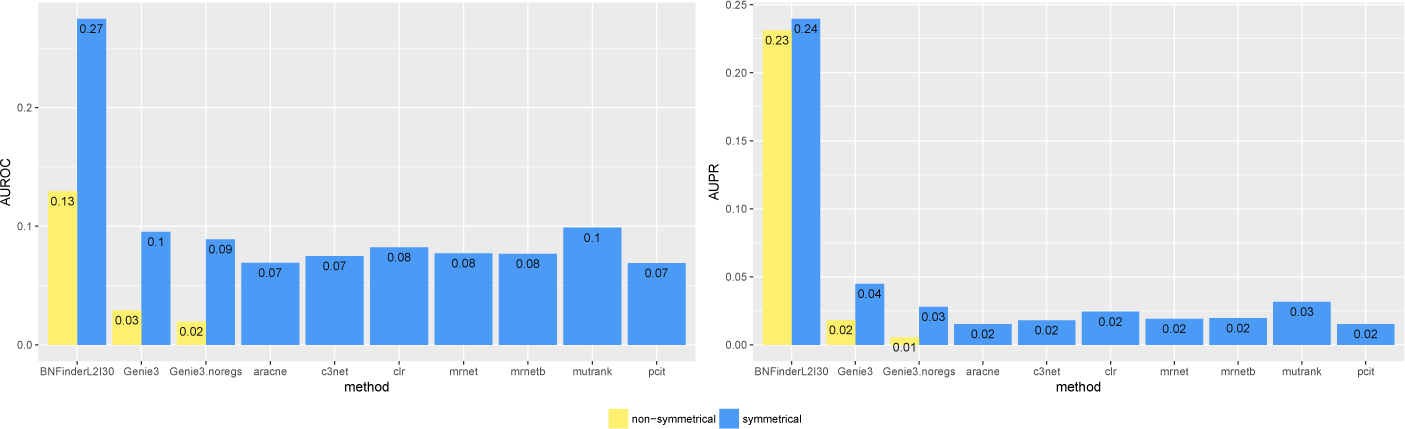
Brem et. al Yeast dataset, evaluation by NetBenchmark package. Directed and non-directed (with symmetrical adjacency matrix) gold standard network evaluation is shown in different colours. Genie3.noregs is the result of Genie3 execution without the regulators list. BNFinder is used with parents sets limit 2 and suboptimal parents sets 30.

We also studied the effect of the number of experiments on the accuracy of inferred network. For GNW2000 synthetic Yeast data we performed two separate tests: one with full dataset - 2000 experiments, and second with only 150 randomly selected observational points. Figure 12 clearly shows that all the methods improved their results on the full dataset, with BNFinder being among the top methods and having best AUPR on the reduced dataset. Interestingly, we did not see such a major difference between BNFinder and other methods in contrast to Brem at al. data, given the same input parameters and both datasets being of Yeast origin. It shows the importance of developing new gold standards based on experimental data from model organisms as synthetic data only cannot reflect all the complexity of biological interactions.

**Figure 12.**
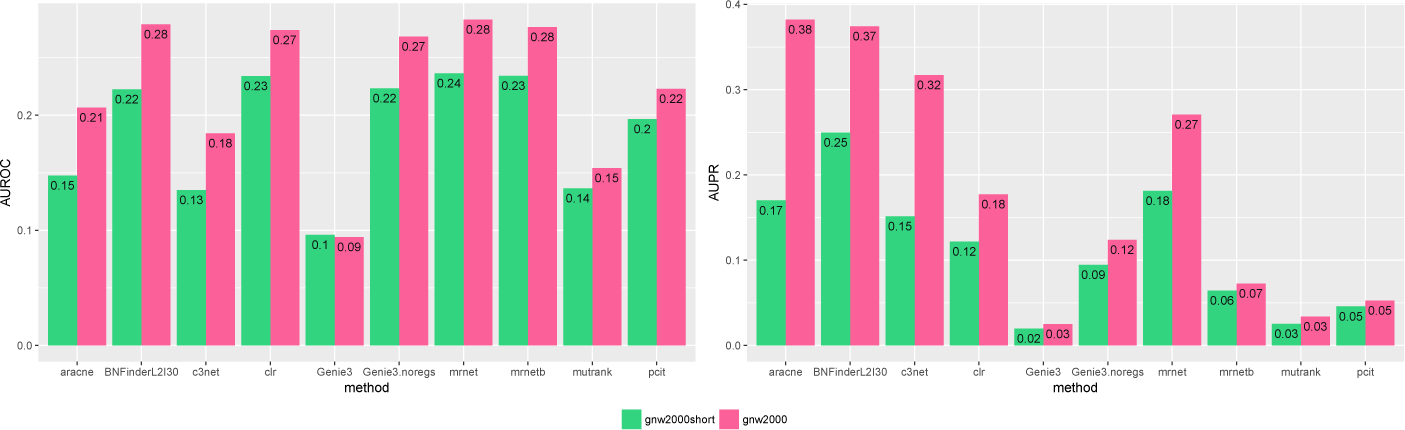
GNW 2000 genes Yeast synthetic dataset, evaluation by NetBenchmark package. GNW 2000 short dataset contains only 150 observations, while GNW 2000 has all the 2000 observations. The effect of observations number increase is clearly seen for all the methods. Genie3.noregs is the result of Genie3 execution without the regulators list. BNFinder is used with parents sets limit 2 and suboptimal parents sets 30.

*Exploring BNFinder parameter space*. The main advantage of BNFinder in comparison with heuristic search Bayesian tools such as Banjo is that BNFinder reconstructs optimal networks, which also means that the same parameters lead to the same result. However, with BNFinder one can use number of input arguments such as scoring functions (Bayesian-Dirichlet equivalence, Minimal Description Length or Mutual information test), static or dynamic (also with self-regulatory loops) BNs, perturbation data, or even prior information on the network structure. All of these may alter results significantly, so, naturally we are interested in choosing best parameters for a particular dataset. Here we studied the impact of two very important parameters: parents sets limit and number of suboptimal parents sets (gives alternative sets of regulators with lower scores).

In Table 4 we have summarized the total number of interactions returned by BNFinder with different maximal parents per gene and different number of suboptimal parent sets. The results indicate that increasing the size of the allowed parent set leads to the decrease in the total returned edges in the network. This may seem surprising at first, but it is consistent with highly overlapping suboptimal parents sets. Theoretically, increasing parents set limit should lead to better precision, while increasing the number of suboptimal parents set should increase the number of false positives by adding lower scored parents. However, it may depend on the particular dataset, especially on the number of observations available. On top of that, in cases where using higher number of parents per variable is computationally challenging, suboptimal parents may compensate for this limitation.

**Table 4.** The number of the interactions in the output network based on different BNFinder parameters. DREAM2 Genome Scale Network Inference data is used.

|  | Subparents=25 | Subparents=50 | Subparents=100 |
| --- | --- | --- | --- |
| Parent limit=1 | 78366 | 156685 | 313061 |
| Parent limit=2 | 65108 | 116424 | 213389 |

We studied the effect of different parameters by plotting AUPR against AUROC values. Figure 13 shows that on 2000 genes x 2000 experiments synthetic dataset using two parents per gene is always better than one, and increasing the number of suboptimal parents leads to increase in AUROC and slight decrease in AUPR values. Inferring a network from the same dataset with only 150 experiments sometimes resulted in lower AUPR for two parents per gene cases in comparison with singleton parents sets. While BNFinder scoring function penalizes for parents set size increase it might still produce false positives when the number of observational points is low and number of variables is more than ten folds bigger. Importantly, zero (or very low) number of sub-optimal parent sets leads to extremely sparse output network (the number of edges is much less than 20% of all possible interaction) and therefore poor AUPR and AUROC values.

**Figure 13.**
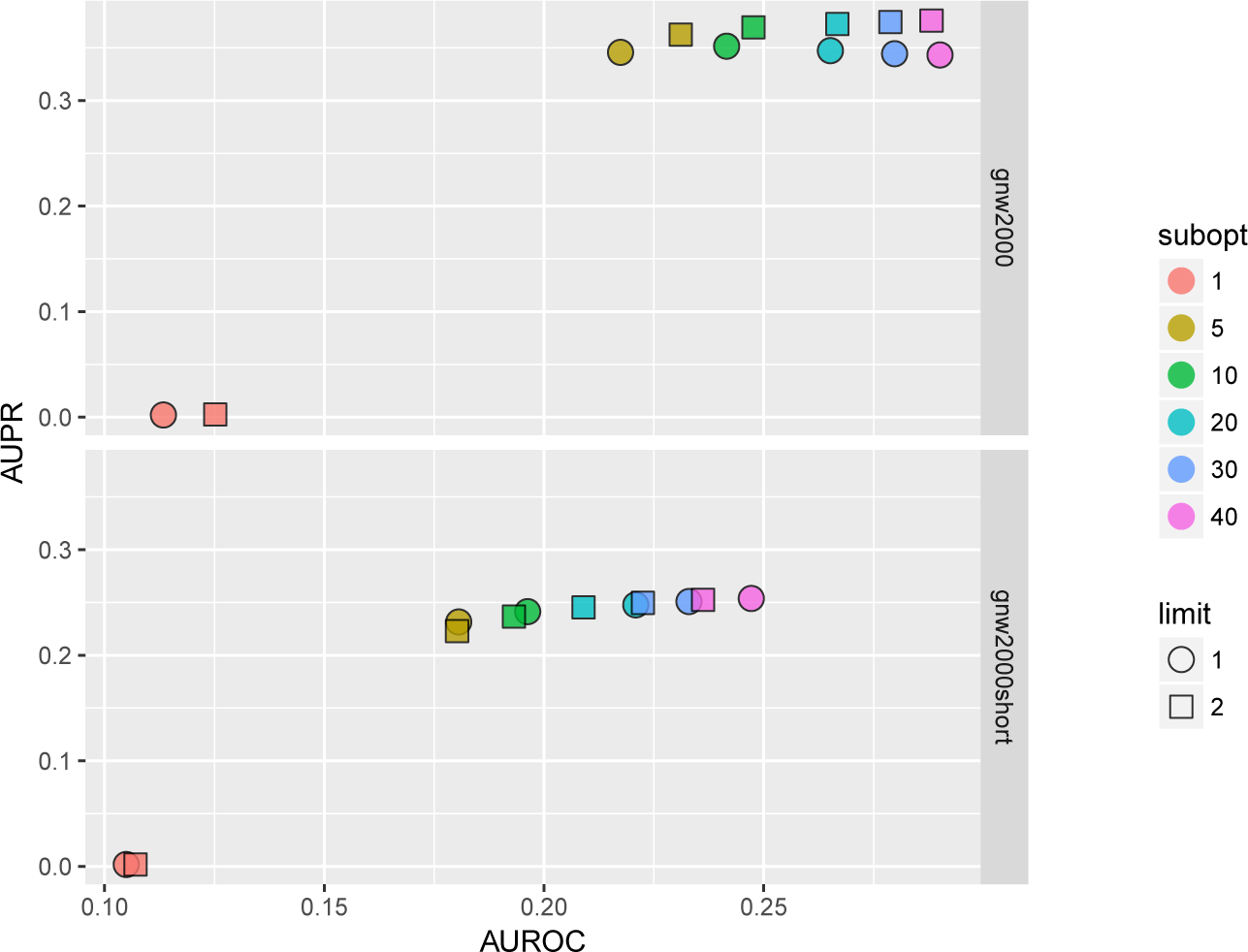
BNFinder parameter space examination on GNW 2000 genes Yeast synthetic dataset, evaluation by NetBenchmark package. GNW 2000 short dataset contains only 150
observations, while GNW 2000 has all the 2000 observations. *Limit* stands for parents set size limit, while *subopt* denotes the number of suboptimal parents set in the output network.

In general we can conclude that if the user is interested in the very top of the strongest interactions in the network, he or she should use small numbers of sub-optimal parent (up to 5) sets and small limit on the parent-set size (up to 3). However, if one is interested in discovering the more global picture of the true regulatory network, one should focus on the higher number of sub-optimal parent sets with limit on the set size as high as it is computationally feasible.

The results of all performance and accuracy tests are available from dedicated github repository - https://github.com/sysbio-vo/article-bnf-suppl.

## Discussion

### Performance

Despite seemingly complex behavior of **hybrid** algorithm and many cases where **hybrid** and **set-wise** algorithms can be applied, we can give the users the best practice guidance for BNFinder application. In case of small networks where number of variables is 2 or more times less than number of cores it is advisable to use **hybrid** algorithm. The same is also applicable when the user imposes parent set limit equal or less than 3, which makes the computational load per variable more even. In case when the complex layered structure of regulators is introduced it is always better to use **set-wise**. And finally, the user may just use the default parameters as **set-wise** algorithm did not show major drop in performance in comparison with **hybrid** one.

### Accuracy of reconstruction

While we understand that there are many more tools for gene regulatory networks reconstruction in the literature we believe that Net-Benchmark package is representative for the field since it incorporates state-of-the-art methods, which are based on variety of different algorithms. On top of that using benchmarking tool makes it easier for other researchers to compare their methods to our results.

Measuring AUROC and AUPR values on 14 different datasets revealed that studied methods behave differently on different datasets, and none of the methods scored best in all cases. In general time series data proved to be more challenging for the methods than inferring network from static gene expression datasets. Our results on 10 genes networks evaluation with top 20% and 100% interactions showed that such small networks can hardly be used as the only source of comparison.

Testing BNFinder on mentioned datasets we concluded that it performed best on the static gene expression datasets with additional prior knowledge (transcription factors list, prior edge probability between genes), while for other methods such as Genie3 the same information did not yield significant improvement.

## Conclusions

Improvement over previous version of BNFinder made it feasible to analyze datasets that were impossible to analyze before by utilizing the power of distributed and parallel computing. It allowed us to significantly extend the application range of the tool and for the first time compare it with best-performing non-Bayesian methods. BNFinder showed overall comparable performance on synthetic and real biological data, providing significant advantage in cases when prior knowledge on genes interactions can be introduced. This can lead to further research on the optimization of the BNFinder method for the purpose of finding larger networks with better accuracy. We provide the new BNFinder implementation freely for all interested researchers under a GNU GPL 2.0 license.

## Availability and requirements

**Project name:** BNFinder

**Project home page:** https://github.com/sysbio-vo/bnfinder

**Source code release used in the article:** https://github.com/sysbio-vo/bnfinder/releases/tag/v2.2

**Supplemental material:** https://github.com/sysbio-vo/article-bnf-suppl

**Operating system(s):** Platform independent

**Programming language:** Python

**Other requirements:** Python 2.4 or higher. Python 3 is not supported

**License:** GNU GPL Library version 2

**Any restrictions to use by non-academics:** None

## Author’s contributions

All the computational experiments, the implementation and majority of text writing were done by AF. BW contributed the initial idea of parallelizing BNFinder, supervised the study and helped with finishing the manuscript.

## Acknowledgements

This work was partially supported by the National Center for Science grant (decision number DEC-2012/05/B/N22/0567) and Foundation for Polish Science within the SKILLS programme. Also this work was supported by National program of Grid technologies implementation and usage in Ukraine (project number 69-53/13 and 0117U002812).

For distributed calculations authors used computational clusters of Taras Shevchenko National University of Kyiv, Institute of Molecular Biology and Genetics of NASU, Institute of Food Biotechnology and Genomics of NASU, joint cluster of SSI “Institute for Single Crystals” and Institute for Scintillation Materials of NASU.

We would like to express out gratitude to Prof. Maria Obolenska from Institute of Molecular Biology and Genetics, Kyiv, Ukraine for the proofreading and critique.

We are extremely grateful to Pau Bellot from Centre for Research in Agricultural Genomics (CRAG), Barcelona, Spain, one of the NetBenchmark R package authors and maintainers, for the assistance with package related problems. Together we were able to fix two bags, which under specific circumstances might have hindered evaluation results. The virtue of Open Source has yet again proven to positively influence scientific advances.

## Additional Files

Additional file 1 – BNFinder source code

BNFinder is written on python, so you will need python 2.4 or higher in order to run it.

## References

1. Friedman, N., Koller, D.: Being bayesian about network structure. a bayesian approach to structure discovery in bayesian networks. Machine learning 50(1-2), 95–125 (2003)

2. Chickering, D.M., Heckerman, D., Meek, C.: Large-sample learning of bayesian networks is np-hard. The Journal of Machine Learning Research 5, 1287–1330 (2004)

3. Dojer, N.: Learning bayesian networks does not have to be np-hard. In: Královic, R., Urzyczyn, P. (eds.) Mathematical Foundations of Computer Science 2006: 31st International Symposium, MFCS 2006, Stará Lesná, Slovakia, August 28-September 1, 2006, Proceedings. LNCS sublibrary: Theoretical computer science and general issues, pp. 305–314. Springer, Berlin/Heidelberg (2006)

4. Wilczyński, B., Dojer, N.: Bnfinder: exact and efficient method for learning bayesian networks. Bioinformatics 25(2), 286–287 (2009)

5. Zou, M., Conzen, S.D.: A new dynamic bayesian network (dbn) approach for identifying gene regulatory networks from time course microarray data. Bioinformatics 21(1), 71–79 (2005)

6. Jansen, R., Yu, H., Greenbaum, D., Kluger, Y., Krogan, N.J., Chung, S., Emili, A., Snyder, M., Greenblatt, J.F., Gerstein, M.: A bayesian networks approach for predicting protein-protein interactions from genomic data. Science 302(5644), 449–453 (2003)

7. Dabrowski, M., Dojer, N., Zawadzka, M., Mieczkowski, J., Kaminska, B.: Comparative analysis of cis-regulation following stroke and seizures in subspaces of conserved eigensystems. BMC systems biology 4(1), 86 (2010)

8. Bonn, S., Zinzen, R.P., Girardot, C., Gustafson, E.H., Perez-Gonzalez, A., Delhomme, N., Ghavi-Helm, Y., Wilczyński, B., Riddell, A., Furlong, E.E.: Tissue-specific analysis of chromatin state identifies temporal signatures of enhancer activity during embryonic development. Nature genetics 44(2), 148–156 (2012)

9. Wilczynski, B., Liu, Y.-H., Yeo, Z.X., Furlong, E.E.: Predicting spatial and temporal gene expression using an integrative model of transcription factor occupancy and chromatin state. PLoS computational biology 8(12), 1002798 (2012)

10. Dojer, N., Bednarz, P., Podsiadło, A., Wilczyński, B.: Bnfinder2: Faster bayesian network learning and bayesian classification. Bioinformatics, 323 (2013)

11. Barabasi, A.-L., Oltvai, Z.N.: Network biology: understanding the cell’s functional organization. Nature Reviews Genetics 5(2), 101–113 (2004)

12. McCool, M., Reinders, J., Robison, A.: Structured Parallel Programming: Patterns for Efficient Computation. Elsevier, Waltham, MA (2012)

13. Sachs, K., Perez, O., Pe’er, D., Lauffenburger, D.A., Nolan, G.P.: Causal protein-signaling networks derived from multiparameter single-cell data. Science 308(5721), 523–529 (2005)

14. Stolovitzky, G., Prill, R.J., Califano, A.: Lessons from the dream2 challenges. Annals of the New York Academy of Sciences 1158(1), 159–195 (2009)

15. DREAM2, Challenge 5 Synopsis. https://www.synapse.org/#!Synapse:syn3034894/wiki/74418

16. Faith, J.J., Hayete, B., Thaden, J.T., Mogno, I., Wierzbowski, J., Cottarel, G., Kasif, S., Collins, J.J., Gardner, T.S.: Large-scale mapping and validation of escherichia coli transcriptional regulation from a compendium of expression profiles. PLoS biology 5(1), 8 (2007)

17. Watkinson, J., Liang, K.-c., Wang, X., Zheng, T., Anastassiou, D.: Inference of regulatory gene interactions from expression data using three-way mutual information. Annals of the New York Academy of Sciences 1158(1), 302–313 (2009)

18. Reich, M., Liefeld, T., Gould, J., Lerner, J., Tamayo, P., Mesirov, J.P.: Genepattern 2.0. Nature genetics 38(5), 500–501 (2006)

19. Ukrainian National Grid Infrastructure. http://infrastructure.kiev.ua/en/

20. Ellert, M., Grønager, M., Konstantinov, A., Kónya, B., Lindemann, J., Livenson, I., Nielsen, J.L., Niinimäki, M., Smirnova, O., Wääanänen, A.: Advanced resource connector middleware for lightweight computational grids. Future Generation computer systems 23(2), 219–240 (2007)

21. Stolovitzky, G., Monroe, D., Califano, A.: Dialogue on reverse-engineering assessment and methods. Annals of the New York Academy of Sciences 1115(1), 1–22 (2007)

22. Yeung, K.Y., Dombek, K.M., Lo, K., Mittler, J.E., Zhu, J., Schadt, E.E., Bumgarner, R.E., Raftery, A.E.: Construction of regulatory networks using expression time-series data of a genotyped population. Proceedings of the National Academy of Sciences 108(48), 19436–19441 (2011)

23. Brem, R.B., Kruglyak, L.: The landscape of genetic complexity across 5,700 gene expression traits in yeast. Proceedings of the National Academy of Sciences of the United States of America 102(5), 1572–1577 (2005)

24. Schaffter, T., Marbach, D., Floreano, D.: Genenetweaver: in silico benchmark generation and performance profiling of network inference methods. Bioinformatics 27(16), 2263–2270 (2011)

25. Bellot, P., Olsen, C., Salembier, P., Oliveras-Vergés, A., Meyer, P.E.: Netbenchmark: a bioconductor package for reproducible benchmarks of gene regulatory network inference. BMC bioinformatics 16(1), 312 (2015)

26. Young, W.C., Raftery, A.E., Yeung, K.Y.: Fast bayesian inference for gene regulatory networks using scanbma. BMC systems biology 8(1), 47 (2014)

27. Meyer, P.E., Lafitte, F., Bontempi, G.: Minet: an open source r/bioconductor package for mutual information based network inference. BMC bioinformatics 9(article 461) (2008)

28. Obayashi, T., Kinoshita, K.: Rank of correlation coefficient as a comparable measure for biological significance of gene coexpression. DNA research 16(5), 249–260 (2009)

29. Margolin, A.A., Nemenman, I., Basso, K., Wiggins, C., Stolovitzky, G., Dalla Favera, R., Califano, A.: Aracne: an algorithm for the reconstruction of gene regulatory networks in a mammalian cellular context. In: BMC Bioinformatics, vol. 7, p. 7 (2006). BioMed Central

30. Reverter, A., Chan, E.K.: Combining partial correlation and an information theory approach to the reversed engineering of gene co-expression networks. Bioinformatics 24(21), 2491–2497 (2008)

31. Altay, G., Emmert-Streib, F.: Inferring the conservative causal core of gene regulatory networks. BMC systems biology 4(1), 132 (2010)

32. Meyer, P.E., Kontos, K., Lafitte, F., Bontempi, G.: Information-theoretic inference of large transcriptional regulatory networks. EURASIP journal on bioinformatics and systems biology 2007, 8–8 (2007)

33. Meyer, P., Marbach, D., Roy, S., Kellis, M.: Information-theoretic inference of gene networks using backward elimination. In: BioComp, pp. 700–705 (2010)

34. Irrthum, A., Wehenkel, L., Geurts, P., et al.: Inferring regulatory networks from expression data using tree-based methods. PloS one 5(9), 12776 (2010)

35. Hung, L.-H., Shi, K., Wu, M., Young, W.C., Raftery, A.E., Yeung, K.Y.: fastbma: scalable network inference and transitive reduction. GigaScience (2017)

36. Kauffmann, A., Gentleman, R., Huber, W.: arrayqualitymetrics—a bioconductor package for quality assessment of microarray data. Bioinformatics 25(3), 415–416 (2009)

37. Bolstad, B.M., Irizarry, R.A., Åstrand, M., Speed, T.P.: A comparison of normalization methods for high density oligonucleotide array data based on variance and bias. Bioinformatics 19(2), 185–193 (2003)

